# Modeling Hormonal Control of Cambium Proliferation

**DOI:** 10.1101/101550

**Authors:** Vladyslav Oles, Alexander Panchenko, Andrei Smertenko

## Abstract

Rise of atmospheric CO_2_ is one of the main causes of global warming. Catastrophic climate change can be avoided by reducing emissions and increasing sequestration of CO_2_. Trees are known to sequester CO_2_ during photosynthesis, and then store it as wood biomass. Thus, breeding of trees with higher wood yield would mitigate global warming as well as augment production of renewable construction materials, energy, and industrial feedstock. Wood is made of cellulose-rich xylem cells produced through proliferation of a specialized stem cell niche called cambium. Importance of cambium in xylem cells production makes it an ideal target for the tree breeding programs; however our knowledge about control of cambium proliferation remains limited. The morphology and regulation of cambium differs from stem cell niches that control axial growth. For this reason, translating the knowledge about axial growth to radial growth has limited use. Furthermore, genetic approaches cannot be easily applied because overlaying tissues conceal cambium from direct observation and complicate identification of mutants. To overcome the paucity of experimental tools in cambium biology, we constructed a Boolean network CARENET (CAmbium Regulation gene NETwork) for modelling cambium activity, which includes the key transcription factors WOX4 and HD-ZIP III as well as their potential regulators. Our simulations revealed that: (1) auxin, cytokinin, gibberellin, and brassinosteroids act cooperatively in promoting transcription of *WOX4* and *HD-ZIP III*; (2) auxin and cytokinin pathways negatively regulate each other; (3) hormonal pathways act redundantly in sustaining cambium activity; (4) individual cells in the stem cell niches can have diverse molecular identities. CARENET can be extended to include components of other signalling pathways and be integrated with models of xylem and phloem differentiation. Such extended models would facilitate breeding trees with higher wood yield.

## Introduction

Competition for solar energy drives axial growth in many plant species resulting in stem elongation. However, longer stems become vulnerable to the forces of gravity and wind. Furthermore, efficiency of photosynthesis in leaves depends on the long-range transport of water and minerals from roots to shoots through the stem. Photoassimilates produced in leaves have to be transported back to roots. Hence, longer stems impose constraints on photosynthesis and growth. In the course of evolution, plants developed mechanisms that balance axial growth with radial (secondary) growth. One of the main outcomes of radial growth is formation of xylem and phloem. Hollow xylem cells reinforced with thick secondary cell walls provide mechanical strength to the stem and a means of root-to-shoot transport. Phloem is responsible for shoot-to-root transport. A specialized meristem located between xylem and phloem, the cambium, divides periclinally to produce precursors for the xylem and phloem cells [1]. Despite significant progress in understanding molecular mechanisms of xylem differentiation and secondary cell wall synthesis, our knowledge about regulation of cambium activity remains scant.

The location of cambium between xylem and phloem facilitates integration of hormonal signals that are produced by both roots and shoots and their translation into secondary growth [2]. Consequently, activity of cambium is known to be regulated non-cell-autonomously by a complex signaling network [3] under control of auxin [4], cytokinin [5], ethylene [6], gibberellins [7], brassinosteroids [8], strigolactones [9] as well as signaling peptides CLE41, CLE42, CLE44 collectively called Tracheary Elements Differentiation Inhibition Factor (TDIF) [10, 11]. However, it remains unclear how complex hormonal signals are integrated by the genetic network that controls cambium.

The best studied signaling module controlling cambium proliferation consistes of TDIF receptor, Phloem Intercalated with Xylem (PXY), a Leucine Rich Repeat domain Receptor-Like Kinase (LRR-RLK) [10, 12], and transcription factors WOX4, WOX14 [13–15] and HD-ZIPIII (represented by ATHB-8 in *Arabidopsis*) [16–18]. While *PXY*, *WOX4*, and *WOX14* are cambium-specific genes, *HD-ZIPIII* is expressed in both cambium and immature xylem cells. TDIF is synthesized in phloem and then diffuses through the apoplastic space to cambium cells where it binds to the receptor domain of PXY [10, 19]. Activated PXY promotes expression of *WOX4*, *WOX14* [13–15], and *ATHB-8* [20]. PXY also phosphorylates and activates GSK3 kinase BIN2 [21]. Phloem is recalcitrant to TDIF signal because transcription of *PXY* in this tissue is inhibited by KANADI [22].

Although numerous lines of evidence support the essential role of PXY/WOX4 signaling module in the regulation of cambium proliferation [23], many questions remain unanswered. First, how is the TDIF signal integrated with other signals? For example, it has been shown that PXY/WOX4 module is also controlled by ethylene [24] and auxin [15]. Second, how are auxin and ethylene signaling integrated with gibberellic acid and brassinosteroids pathways which also control activity of cambium [7, 8]? Third, what makes cambium dormant during the cold season or inactive towards the end of developmental cycle in annual and perennial species?

Understanding the biology of cambium remains incomplete because cambium is concealed under phloem, epidermis, and cortex tissues. This hinders identification of mutants with altered secondary growth and identification of genes that control cambium activity. Furthermore, isolation of live cambium cells has not been achieved thus far. Mathematical modelling can help in predicting the outcome of interactions between components of complex genetic networks. For example, models have been developed for understanding identity of tissues in *Arabidopsis* vascular bundles [20] or for differentiation of xylem cells [25, 26]. However, a model of cambium proliferation has not been created thus far.

Here we describe a network model composed of known regulators of procambium or cambium development and activity, which we call CARENET (CAmbium REgulation gene NETwork). CARENET is a Boolean network of the type originally introduced by Kauffman [27–30]. Such models can be constructed on the basis of mostly qualitative information concerning the cause-and-effect relationships between pairs of agents (e.g. gene A activates or inhibits gene B). Since this type of information is commonly available in the biological literature, Boolean models have an advantage over other types of models (e.g. Ordinary Differential Equations; ODEs) construction of which may require relatively hard to obtain information about reaction rates. Additional information on the analysis and simulation of Boolean networks in biology can be found in the books by Shmulevich and Dougherty [31,32] and in the papers [33,34].

In this work CARENET is primarily used for unraveling interactions between different hormonal signaling pathways for the control of secondary growth. Our simulation experiments accurately represent experimental data on the importance of cytokinin, auxin, and ethylene for cambium activity and demonstrate the ability of gibberellic acid and brassinosteroids to increase activity of cambium. Our model can be used for designing plants with altered secondary growth and biomass yield.

## Materials and Methods

### Software implementation

All simulations of cambium cell were performed using software designed specifically for this study. The program was implemented using Python 2.7 language. It embeds theoretically developed update rules for the model (S1 Table). Primary functionality of the program is to simulate the evolution of production of relevant chemicals in a cambium cell.

The software incorporates a feature that allows application of so-called “control actions”, which manually override the state of any chosen node at any time step. This feature allows simulation of the effect of gene knockout. For instance, control actions forcing PXY node into the state of 0 at every time step of simulation simulates *pxy* mutant.

The program can process multiple initial states of the model in bulk, automatically calculating statistical data for each of the final states found. A single run tests all possible initial states for a specific configuration. Although such automation speeds up the process, simulations are computationally intensive due to the model size: processing of one control configuration required about two hours on a desktop computer.

### Chi-squared test of independence

In order to test statistical significance of the relationships between intracellular hormone accumulation (activity) and proliferation activity (defined in Section “Numerical experiments and their statistical analysis”), or accumulation of another hormone, we use a test of independence, which is a version of the Pearson′s chi-squared test. Since activity is a continuous variable taking values between 0 and 1, and the test only applies to categorical data, we bin this range into several equal parts, e.g., [0,0.25], [0.25,0.5], [0.5,0.75], [0.75,1]. For the control nodes categorization is straightforward, as they can only take values 0 or 1.

Once the values of both variables under consideration are categorized, a contingency table matching categories of one variable with categories of another is constructed. Each cell of the table corresponds to a pair of categories and stores the number of instances for which the variables belong to the respective categories. The null hypothesis states the variables are independent, hence the expected numbers are equal in each cell. Afterwards, the sum of normalized squared deviations of observed numbers from the expected numbers is calculated to obtain the chi-squared test statistic (χ^2^) Its critical value is determined by the degrees of freedom and chosen significance level, which represents the cut-off value for the likelihood of obtaining the experimental result by chance. We set the significance level to 0.05 (5%), commonly accepted as the optimal value. Comparing χ^2^ to the critical value tells us whether we should accept or reject the null hypothesis.

A conclusion to reject the null hypothesis means that two variables are related. In such a case, we estimate whether their correlation is positive or negative. The distributions of activity levels of one variable are compared amongst themselves as another variable changes its activity level from lower to higher. If the first variable gets fewer cases of low activity and more cases of high activity as the second variable increases its level, the correlation is considered to be positive. Conversely, the correlation is considered negative if the above occurs for lower activity of the second variable.

## Results

### Constructing a model for the cambium-regulating gene network

To model cambium activity in response to developmental cues we choose a Boolean approach because it has been shown to produce reliable models of complex interactions even in the absence of quantitative data on concentration of relevant chemicals [34–36]. Here, cambium proliferation is modeled by a deterministic Boolean network (or graph) consisting of nodes and edges (Fig 1). Selection of the nodes and edges is based on information collected from published data on diverse developmental processes including leaf venation, primary growth of roots and stems, and secondary growth of stems and cotyledons in *Arabidopsis*, poplar, and peas (S2 Table). While nodes represent hormones, proteins, genes or cellular processes, the edges represent functional interactions between the nodes. An edge directed from node *A* to node *B* represents a cause and effect relationship between these nodes whereby node *A* either activates node *B* or inhibits it. Thus, each edge in our model is designated as inhibitory or activatory.

**Fig 1.**
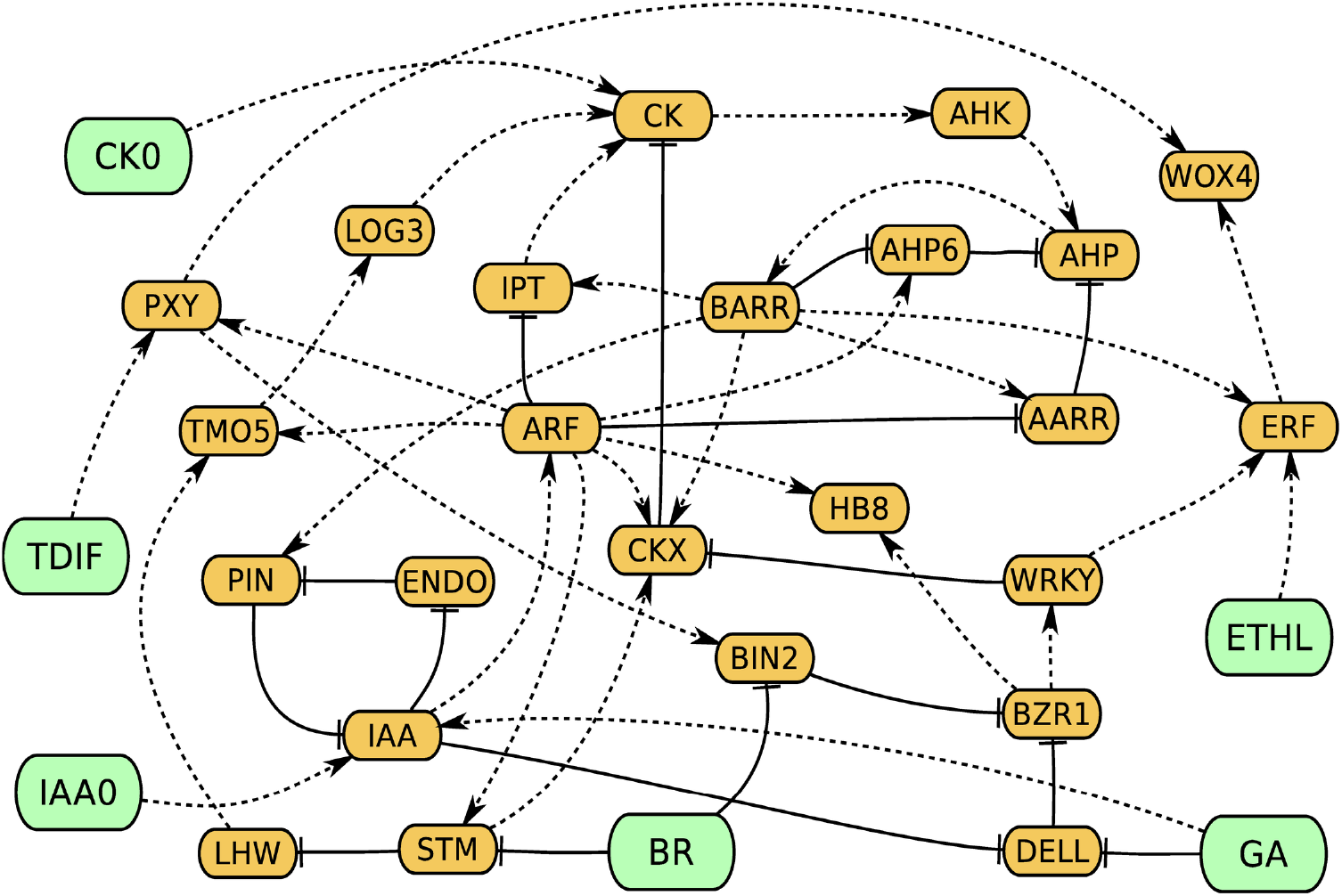
Cambium regulation gene network (CARENET). Control nodes are in green and internal nodes are in orange. Dashed and solid lines indicate activation and inhibition respectively. ATHB8 is labelled as HB8, ENDO means endocytosis.

Inclusion of nodes and edges inferred from research on divergent systems is justified by recent studies which demonstrate similarity of molecular mechanisms underlying cambium activity. For example, PXY-dependent signaling promotes cambium in *Arabidopsis* and poplar [14, 19, 37]. Below we describe the main signaling blocks of the CARENET and justification for inclusion of specific nodes and edges.

#### Cytokinin (CK)

Mutations in genes encoding cytokinin synthesis enzymes (e.g. isopentenyltransferase; IPT) or receptors of cytokinin (Arabidopsis Histidine Kinase; AHK) exhibit reduced xylem content [5, 38–40]. Cytokinin can be synthesized locally and work in a cell-autonomous manner [41] or be transported for long distances through symplastic connections in phloem and act in a non-cell-autonomous fashion [42]. We introduced external (CK0) and internal (CK) nodes to represent both sources of cytokinin. Intracellular homeostasis of cytokinin is maintained through the balance of biosynthesis by IPT and cytokinin nucleoside 5-monophosphate phosphoribohydrolase (LOG), and deactivation by cytokinin oxidase (CKX). The downstream signaling processes contain a self-inhibitory mechanism: CKX transcription is up-regulated through B-type response regulator (RRB) ARR2 [43]. In addition to these components our network also includes AHP, type-A response regulator (RRA), and inhibitor of cytokinin signaling AHP6 (S2 Table). The heterodimer of transcription factors TMP5-LHW contributes to the cytokinin signaling network by promoting cytokinin synthesis through transcriptional activation of LOG3/4 [44].

#### Auxin (IAA)

Auxin transported from shoots and secreted by the surrounding cells (IAA0) is taken up by cambium cells (IAA). The concentration of auxin was found to be the highest in cambium zone of poplar stems [4]. Decapitation of *Arabidopsis* plants shuts down the auxin supply from the shoots and reduces cambium activity while application of exogenous auxin reconstitutes cambium activity [45]. Furthermore, mutants with reduced sensitivity to auxin (e.g. *auxin resistant 1*), exhibit lower activity of interfascicular cambium [9]. Perception of the auxin signal ultimately results in activation of transcriptional regulators Auxin Response Factors (ARF; reviewed in [46]). As the bulk of auxin is produced in the shoots, transport becomes an essential part of the signaling network. Consequently, plants have evolved several auxin transporters amongst which are efflux carriers PIN proteins [47]. Four members of the PIN gene family *PIN1,5,6*, and *8* express in vasculature and play an important role in vasculature patterning [48, 49]. PINs in the surrounding tissues would promote cambium activity by increasing auxin concentration in cambium cells. However, up-regulation of PIN proteins in cambium cells would promote auxin efflux, thus causing reduction of intracellular auxin concentration (IAA) and inhibition of cambium proliferation. In our model, PINs are included in the negative feedback mechanism of intracellular auxin concentration (IAA).

#### Auxin and cytokinin cross-talk

Auxin and cytokinin reciprocally control their intracellular concentration (IAA and CK) through negative feedback loops. Antagonistic relationships between these hormones defines cambium division rate and the vascular tissue pattern in roots [50, 51]. Auxin can cause a reduction of intracellular cytokinin by promoting expression of a key deactivating enzyme CKX [52] and by down-regulating transcription of an essential biosynthetic enzyme *IPT* [53, 54]. In agreement with these observations, treatment with auxin reduces several major cytokinin intermediates [53]. Furthermore, ARFs promote transcription of *AHP6* which inhibits cytokinin signaling [50, 55].

Cytokinin can potentially diminish availability of auxin to cambium cells through RRB-dependent down-regulation of PIN expression in the surrounding tissues [42, 56]. Moreover, cytokinin can reduce the concentration of auxin in cambium cells by up-regulating expression of PIN proteins [57, 58], which depends on binding of the Cytokinin Response Factors (CRFs) to a specific region of *PIN* promoter [59]. In our model cytokinin dampens auxin signaling through RRB-dependent up-regulation of PINs expression and subsequent increase in auxin efflux. This negative feedback loop maintains hormonal balance in cambium under steady state conditions.

Relationships between auxin and cytokinin can be cooperative. In our model ARF promotes transcription of *TMO5* which together with LHW up-regulates expression of a key cytokinin biosynthesis enzyme *LOG3/4* [51]. At the same time, ARF can inhibit transcription of *LHW* through transcriptional activation of *STM* (Fig 1; [60, 61]). It has also been shown that STM may play a role in xylem cells differentiation [62].

#### Ethylene (ETHL)

Ethylene was implicated in secondary growth because transcription of genes encoding Ethylene-Responsive transcription Factors (ERF) are up-regulated in cambium proliferation mutants *pxy* and *wox4* [24]. In agreement with this suggestion, *ERFs* knockout exacerbates the reduced xylem phenotype in *pxy* and reduces transcription level of *WOX4*. These findings indicate that ethylene signaling cooperates with PXY to control WOX4-dependent proliferation of cambium.

#### Brassinosteroids (BR)

Overexpression of the brassinosteroids receptor BRI1 promotes proliferation of cambium and increases the amount of xylem in vascular bundles [63]. A similar phenotype was observed in plants expressing constitutively active BRI1 [8, 64]. BRI1 was omitted in the model because brassinosteroids signal ultimately inhibits protein kinase activity of BIN2. Consequently, degradation of transcription factors *BRASSINAZOLE RESISTANT 1 (BZR1)* and *bri1*-*EMS*-*SUPPRESSOR 1* (*BES1*) is inhibited and they promote transcription of the targets [65]. Only BZR1 was retained in the network because these transcription factors were considered functionally redundant in regulating cambium proliferation. The connection between brassinosteroids pathway and PXY/WOX4 module remains unknown and we propose transcription factors of WRKY family as a potential bridge. This hypothesis is justified by three independent observations. First, *WRKY12* is down-regulated in the bri1 background [66]. Second, according to the publicly available microarray data, transcription of *WRKY12* is up-regulated by BR. Third, ectopic expression of constitutively active mutant BRI1Y831F up-regulates transcription of *WRKY48* [8]. We also propose a connection between WRKY and PXY/WOX4 module through ERF considering the observation that knockout of *Medicago truncatula WRKY12* homologue leads to down-regulation of *ERF4* [67].

#### Gibberellic acid (GA) and crosstalk between GA, BR, and IAA pathways

Over-production of GA in transgenic poplar expressing GA 20-oxidase, an enzyme responsible for GA-biosynthesis, promotes cambium proliferation [7]. Application of GA to decapitated poplar trees also stimulates proliferation of cambium [68]. The GA-signaling pathway was suggested to promote secondary growth during flowering in Arabidopsis because xylem content was three times lower in GA biosynthesis mutant *ga1-3* [69]. The underlying mechanism of interaction between GA signaling and the PXY/WOX4 module remains unknown. Experimental data suggests that cross-talk of the TDIF/PXY signaling module with brassinosteroids and gibberellic acid could be facilitated by members of the WRKY family of transcription factors. In addition to the arguments given in “Brassinosteroids (BR)” paragraph, transcription of *WRKY12, WRKY48*, and *WRKY53* is up-regulated by brassinosteroids according to publicly available microarray data. Furthermore, *WRKY12* is up-regulated by gibberellic acid. WRKY also links brassinosteroids and gibberellic acid with cytokinin signaling by inhibiting transcription of the cytokinin-degrading enzyme CKX [67] and by promoting the LHW-dependent transcription of a gene that encodes the cytokinin biosynthetic enzyme *LOG3/4* [61,66]. Björklund et al. have shown that GA can activate IAA signaling in cambium by promoting expression of PIN1 in the cells at the early xylem differentiation stages [68]. This would ultimately result in higher IAA concentration in cambium cells.

### Model implementation

The nodes of the network are denoted by *V*_1_, *V*_2_…*V*_*N*_, where *N* = 30 is the number of nodes. The changes in gene expression level or concentration of a chemical are modeled by assigning a state s_i_ to the corresponding node *V*_*i*_, *i* = 1…*N*. At each instant of time, a state *s*_*i*_ equals either 0 or 1. The value 0 indicates low activity level, while the value 1 indicates high activity level.

The nodes are divided into two groups: control nodes and internal nodes (Fig 1). Control nodes represent hormonal/peptidic signals produced by surrounding or remote tissues (CK0, IAA0, BR, GA, TDIF, and ETHL). These nodes are not influenced by other nodes in the network. Internal nodes can be regulated by both the control nodes and other nodes in the network. The states of control nodes remained fixed throughout each simulation run, while the states of internal nodes may vary.

As mentioned in Section “Constructing a model for the cambium-regulating gene network”, functional interactions between the nodes are represented by edges. An edge directed from *V_*i*_* to *V_*j*_* encodes the following assumptions: (i) there is a chain of chemical reactions where *V_*i*_* is an input and *V_*j*_* is a product; (ii) the reactions producing V_*j*_ from V_*i*_ may involve other chemicals but these chemicals are not included into the model; and (iii) these intermediate chemicals do not influence the states of other nodes. An activatory edge from *V_*i*_* to *V_*j*_* is denoted by (*V*_*i*_→*V*_*j*_) and an inhibitory edge is denoted by (*V*_*i*_⊣*V*_*j*_)

### Update rules

The state of each node changes in time according to the update rules. The next value s_*i*_ is determined by the current value *s*_*j*_ of every node affecting *V_*i*_* by means of an edge directed from *V_*j*_* to *V_*i*_*. In order to describe the rules in detail, we use the following notations. Each edge from *V_*j*_* to *V_i_* is assigned a weight

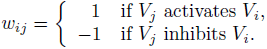

Furthermore we define an auxiliary function 
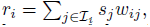
 where 
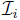
 is the set of indices of all nodes influencing *V_*i*_*. Detailed explanation of the role of *r*_*i*_ in calculating the rules will be provided below. Suppose that there are *n_*i*_* nodes affecting *V_*i*_*, of which *m_*i*_* nodes are inhibitory. As the states *s*_*j*_ of these nodes are either 0 or 1, the range of *r*_*i*_ is the interval [−*m*_i_; *n_i_ −m_*i*_*]. The smallest value −*m*_*i*_ is obtained when the state of every inhibitory node is 1, and the state of every activatory node is 0. The largest value *n*_*i*_ − *m*_*i*_ corresponds to the reverse situation.

To calculate the value of *s*_*i*_ at the next step we compute *r_*i*_* using current values of *s_*j*_* and then apply the following rule:

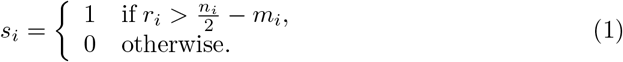

Equation (1) means that *s_*i*_* will take the value 1 if *r_*i*_* is greater than the arithmetic mean of end point values of the interval [−*m*_*i*_, *n*_*i*_ −*m*_*i*_], otherwise *s_i_* will take the value 0.

To illustrate the process of updating the node states using the above rules, we provide an example in which a node *V_*i*_* is influenced by nodes *V*_1_, *V*_2_ and *V*_3_. Suppose that *V*_1_ and *V*_2_ activate *V_i_*, while *V*_3_ inhibits it (Fig 2A). Therefore, there are three edge weights *w*_*i*1_ = 1, *w*_*i*2_ = 1, and *w*_i3_ = − 1, and thus *n_i_* = 3 and *m*_*i*_ = 1. The next value of *s*_*i*_ calculated from the current values of *s*_1_, *s*_2_ and *s*_*3*_ according to (1) is shown in Fig 2B.

**Fig 2.**
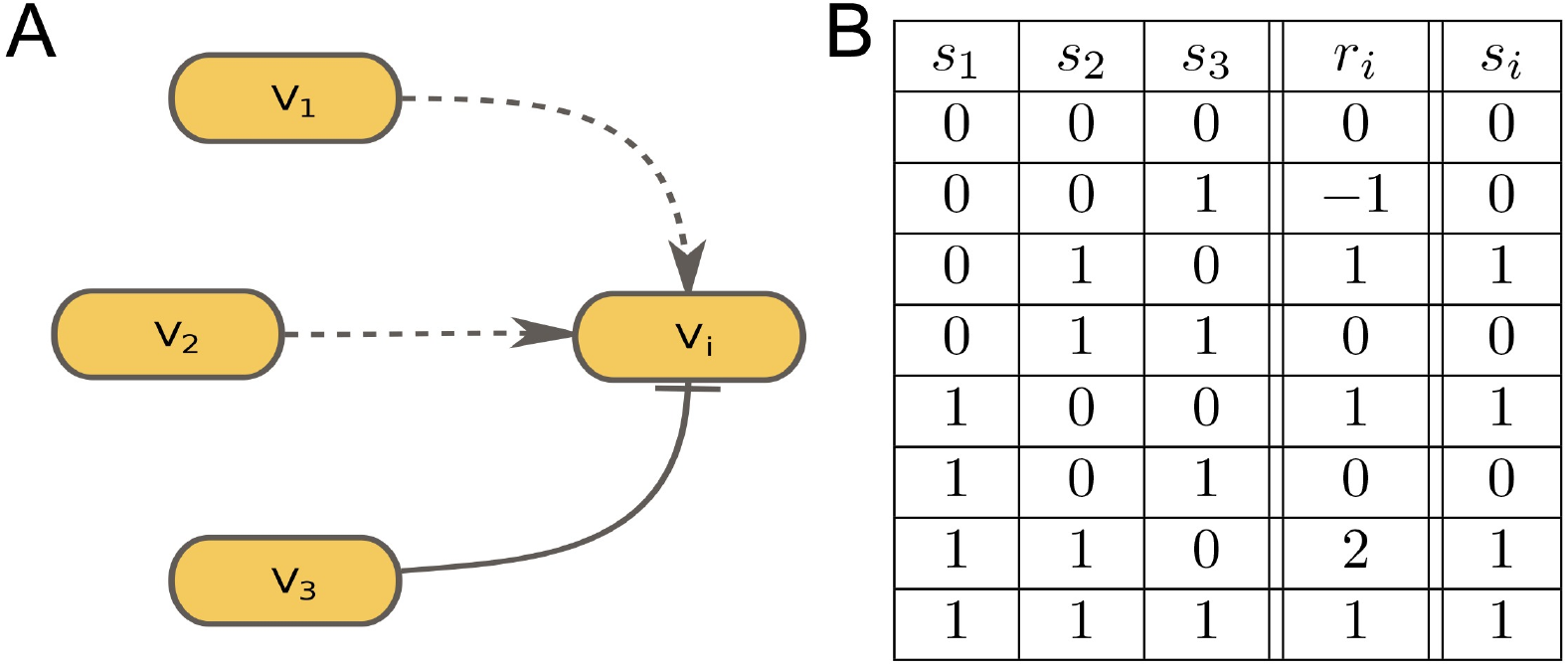
Interaction rules between the nodes. A, Example of a node *V_i_* influencing nodes *V*_1_, *V*_2_ and *V*_3_. B, Values of *r_i_* and *s_i_* for all possible combinations of *s*_1_, *s*_2_ and *s*_3_.

#### Remark on why the inequality in (1) is strict

When 
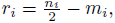
 the value of *s_*i*_* obtained from the formula (1) equals 0. This choice is motivated by the special case in which a node *V_*i*_* is influenced by two nodes connected by activatory edges. When one of them is active and the other is inactive, the resulting 
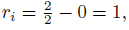
 which is the midpoint of the range interval [0; 2]. Assuming that both nodes must be active in order for *s*_*i*_ = 1, in this case the node *V_*i*_* should be inactive (s_i_ = 0). For the sake of consistency, we adopt the same rule for the generalcase of *n_*i*_* nodes: if 
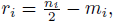
then *s*_*i*_ = 0.

### Numerical experiments and their statistical analysis

Simulating dynamics of genetic networks requires information on the initial conditions. However, limited time resolution of currently available assays of gene transcription and protein turnover rate in cambium precludes generating sufficiently accurate datasets that could be used as initial conditions. Hence, information about behavior of the network is generated by repeatedly simulating its dynamics under all possible initial conditions. A single simulation run consists of the following steps. First, each control node is assigned a specific state. Combination of such assignments for all control nodes constitutes a *control state*. Once chosen, the control state remains fixed throughout the course of the run. Second, initial states are assigned to the internal nodes. Third, the states of all internal nodes are updated according to the rules described above.

The states of the internal nodes are updated sequentially until a *final state* is attained. In this state the network either remains completely static (a steady state), or cycles through a finite number of states (a limit cycle). As the outcome of a run we record the initial conditions (initially assigned states of the internal nodes), the control state, and the final state. Then, the initial conditions and the controls are modified and a new run commences. The simulation stops once all combinations of the initial conditions and all control states have been exhausted. The final states provide reliable information about behavior of the network, while transient states are considered uninformative because Boolean dynamical systems lack physically relevant time scales [20,70].

Each numerical experiment consists of many runs performed with the same values of controls and different initial conditions for the internal nodes. Although the final states obtained for specific initial conditions could potentially be different, many initial conditions lead to the same final state. Hence, the number of final states compatible with a given control state is smaller (typically orders of magnitude smaller) than the number of all possible initial conditions. This suggests that the number of initial conditions leading to a particular final state can be used to characterize its persistence relative to other final states.

The lack of experimental data on the initial conditions of genetic networks is reminiscent of molecular dynamics where the initial positions and velocities of individual molecules can not be measured. Thus, observable quantities in molecular dynamics are produced by means of probabilistic (ensemble) averaging. Every set of initial positions and velocities is assigned a number reflecting the probability of occurrence. Then observed or simulated results corresponding to different initial conditions are averaged using these probabilities as weights.

Here, we develop a similar approach for interpreting dynamic behavior of Boolean networks. Within this approach, each final state *F* is assigned a weight

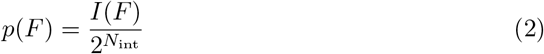

Where *I*(*F*) is the number of initial conditions which lead to *F* and *N_*int*_* is the number of interior nodes. The weight *p*(*F*) takes values between 0 and 1 and the sum of *p*(*F*) over all possible *F* equals 1. Thus *p*(*F*) can be viewed as the probability of occurrence of the final state *F*. It is also worth noting that the assignment (2) is based on the assumption that all initial conditions are equiprobable. Such assignment of probabilities based on the size of attraction basins have been previously employed in [71] for studying the basin entropy of Boolean networks.

The relative importance of the node *V_*i*_* in a final state *F* is described by the *activity α_*i*_*(F)**, taking values between 0 and 1. If *F* is a steady state, *α*_*i*_*(F)* = *s_i_*, so activity is either 0 or 1. If *F* is a limit cycle, a node state *s*_*i*_ may take the value 1 at some steps within the cycle, and take the value 0 at other steps. In this case, *α*_*i*_*(F)* is determined by dividing the number of steps where *s*_*i*_ = 1 by the total length of the cycle. For example, if a limit cycle F consists of 7 update steps with 3 of them resulting in *s*_*i*_ = 1, then 
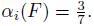

Next, we define the average activity 
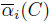
 associated with a control state *C*. It is calculated as a weighted mean of activities *α*_*i*_(*F*_*k*_) taken over all final states **F*_*k*_* related to *C*. Related final states are defined as states that are produced by simulating network dynamics with a fixed control state *C* and all possible initial conditions. Specifically,

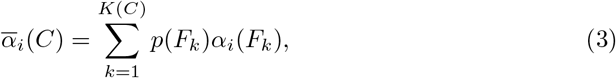

Where *F_*k*_*, *k* = 1…*K*(*C*) denote all final states related to *C*. By combining formulas and (3) it can be easily seen that 
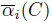
 is the arithmetic mean of the activities *α*_*i*_ resulting from each of 
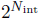
 initial conditions (with the control state *C* being fixed).

### Reporters of the CARENET activity

Available experimental data demonstrates that transcription levels of *WOX4* and *AtHB8* correlate with cambium proliferation and therefore can be used to assess the activity of CARENET. The transcription pattern of *WOX4* is spatially limited to (pro)cambium [14,15,72]. Furthermore, reduced cambium proliferation in *pxy* and ethylene signaling mutants *erf108erf109* is accompanied by lower *WOX4* transcription level [13,24]. Another important argument for using *WOX4* as a reporter of cambium activity is that exogenous application of auxin in wox4 background promoted transcription of *CLE44* (TDIF) peptide; however proliferation of cambium was not induced [15]. *AtHB8* is transcriptionally active in the pre-procambial cells of leaf veins [73] and *AtHB8* transcription also increases around damaged parts of stem [16]. Presence of *AtHB8* in the protoxylem and metaxylem domain of vascular bundles [50] suggests an additional role of AtHB8 in the transition of cambium cells from proliferation to differentiation.

Thus, two parameters can be potentially used to describe quantitatively the propensity of a control state *C* to induce proliferation: one relies on 
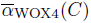
only, and another that takes into account the activities of both nodes WOX4 
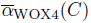
and AtHB8 
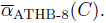
In the latter case, the average combined proliferation activity 
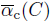
 associated with C is defined as follows:

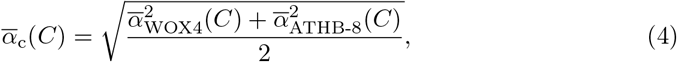

### Overall Performance of the CARENET

Simulation of the CARENET using all possible combinations of control states and initial conditions (2^30^) resulted in 204 final states. Amongst these, 152 states were limit cycles consisting of 2 to 18 steps (S3 Table). Several representative examples of the limit cycles and one steady state are shown in Fig 3 and S1-S3 Figs. Structure of the limit cycles shows periodic oscillations of cytokinin signaling nodes CK, AHK, or AHP irrespectively of the status of external cytokinin (CK0). These oscillations are enabled through the activity of nodes that produce (LOG3 and IPT) or degrade (CKX) cytokinin. Furthermore, statistical analysis of the final states revealed a negative correlation between IAA and CK (Table 1), which was independent of external auxin and cytokinin. These outcomes agree with the experimental data on the mutually inhibitory relationships between cytokinin and auxin signaling.

**Table 1.** Correlation analysis between control nodes and proliferation. Grey shading indicates statistically significant correlation between the variables (0.05 significance level)

| Variable 1 | Variable 2 | Conditions | Relationships |
| --- | --- | --- | --- |
| IAA | CK | – | Negatively correlate with each other |
| IAA | CK | CK0 = 0,<br>IAA0 = 1 | Negatively correlate with each other |
| IAA | CK | CK0 = 1,<br>IAA0 = 1 | Negatively correlate with each other |
| $\bar{\alpha}_c(C)$ | $\bar{\alpha}_{\text{WOX4}}(C)$ | – | $\bar{\alpha}_{\text{WOX4}}(C)$ positively correlates with $\bar{\alpha}_c(C)$ |
| CK | $\bar{\alpha}_c(C)$ | – | CK and are dependent, but the character of relationships could not be determined |
| IAA | $\bar{\alpha}_c(C)$ | – | IAA positively correlates with $\bar{\alpha}_c(C)$ |
| ETHL | $\bar{\alpha}_c(C)$ | – | ETHL positively correlates with $\bar{\alpha}_c(C)$ |
| GA | $\bar{\alpha}_c(C)$ | – | Test is inconclusive at 0.05 significance level |
| BR | $\bar{\alpha}_c(C)$ | – | BR positively correlates with $\bar{\alpha}_c(C)$ |
| TDIF | $\bar{\alpha}_c(C)$ | – | TDIF positively correlates with $\bar{\alpha}_c(C)$ |
| CK | $\bar{\alpha}_c(C)$ | TDIF = 1 | CK positively correlates with $\bar{\alpha}_c(C)$ |
| IAA | $\bar{\alpha}_c(C)$ | TDIF = 1 | IAA positively correlates with $\bar{\alpha}_c(C)$ |
| ETHL | $\bar{\alpha}_c(C)$ | TDIF = 1 | ETHL positively correlates with $\bar{\alpha}_c(C)$ |
| GA | $\bar{\alpha}_c(C)$ | TDIF = 1 | GA positively correlates with $\bar{\alpha}_c(C)$ |
| BR | $\bar{\alpha}_c(C)$ | TDIF = 1 | BR positively correlates with $\bar{\alpha}_c(C)$ |
| CK | $\bar{\alpha}_c(C)$ | – | CK positively correlates with $\bar{\alpha}_{\text{WOX4}}(C)$ |
| IAA | $\bar{\alpha}_{\text{WOX4}}(C)$ | – | Test is inconclusive at 0.05 significance level |
| ETHL | $\bar{\alpha}_{\text{WOX4}}(C)$ | – | ETHL positively correlates with $\bar{\alpha}_{\text{WOX4}}(C)$ |
| GA | $\bar{\alpha}_{\text{WOX4}}(C)$ | – | Test is inconclusive at 0.05 significance level |
| BR | $\bar{\alpha}_{\text{WOX4}}(C)$ | – | Test is inconclusive at 0.05 significance level |
| TDIF | $\bar{\alpha}_{\text{WOX4}}(C)$ | – | TDIF positively correlates with $\bar{\alpha}_{\text{WOX4}}(C)$ |
| CK | $\bar{\alpha}_{\text{WOX4}}(C)$ | TDIF = 1 | CK positively correlates with $\bar{\alpha}_{\text{WOX4}}(C)$ |
| IAA | $\bar{\alpha}_{\text{WOX4}}(C)$ | TDIF = 1 | IAA positively correlates with $\bar{\alpha}_{\text{WOX4}}(C)$ |
| ETHL | $\bar{\alpha}_{\text{WOX4}}(C)$ | TDIF = 1 | ETHL positively correlates with $\bar{\alpha}_{\text{WOX4}}(C)$ |
| GA | $\bar{\alpha}_{\text{WOX4}}(C)$ | TDIF = 1 | GA positively correlates with $\bar{\alpha}_{\text{WOX4}}(C)$ |
| BR | $\bar{\alpha}_{\text{WOX4}}(C)$ | TDIF = 1 | BR positively correlates with $\bar{\alpha}_{\text{WOX4}}(C)$ |

**Fig 3.**
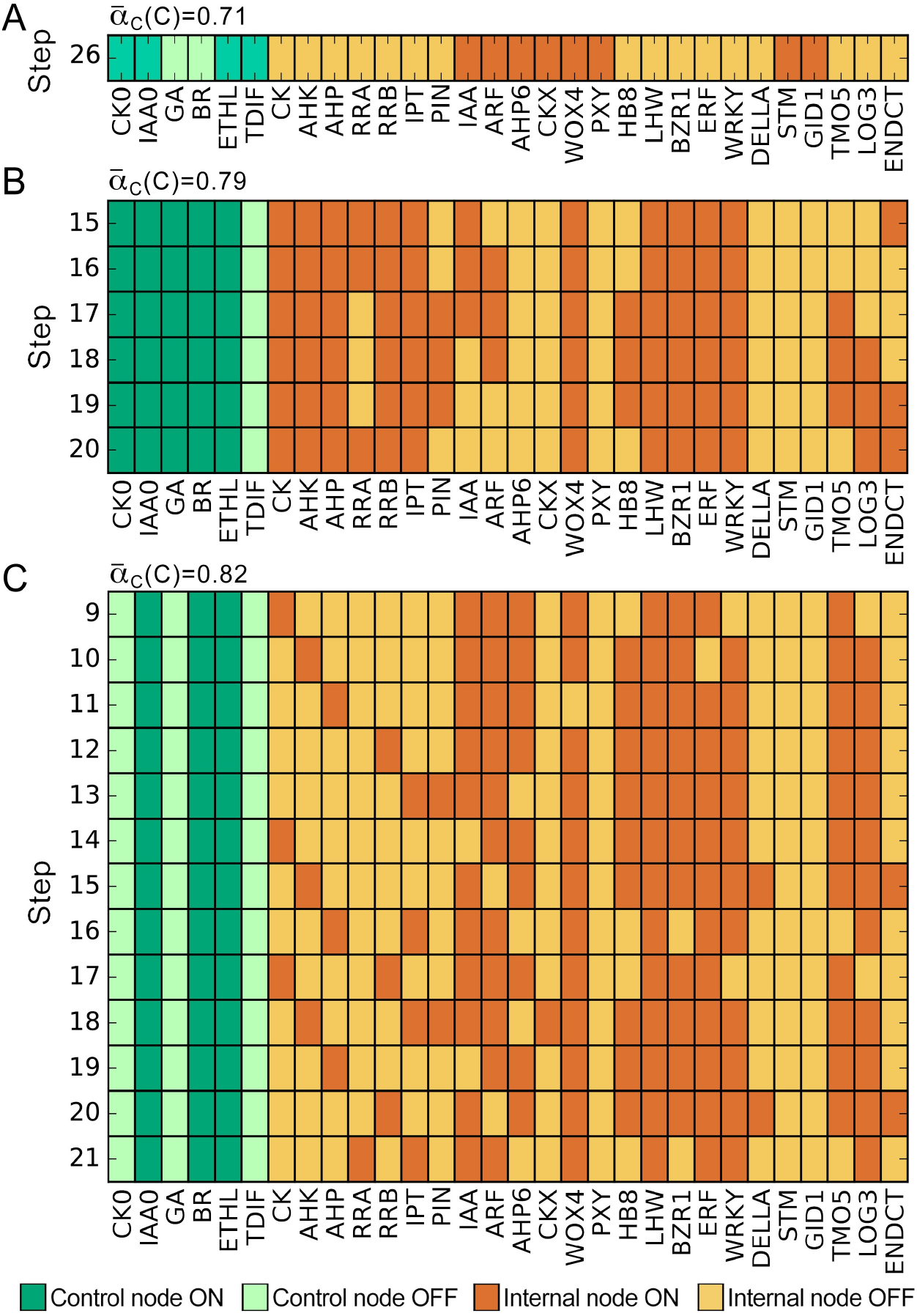
Representative examples of a stable state and limit cycles. A, Stable state achieved by step 26. B, C, Limit cycles established by step 15 or 9 respectively. Intracellular cytokinin (CK) synthesis oscillates even in the absence of external cytokinin CK0. ATHB8 is labelled as HB8.

As described earlier, two parameters can potentially be used for evaluating CARENET behavior: one that relies on the activity of WOX4 node, 
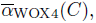
and another one that takes into account activity of WOX4 and AtHB8 nodes, 
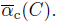
We found that 
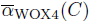
and 
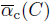
are significantly correlated (Table 1). Therefore, 
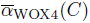
and 
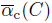
can be used for assessing activity of CARENET interchangeably.

### Regulation of cambium activity by hormones

To examine the responsiveness of the CARENET to external signals, we measured the impact of individual hormones on cambium proliferation (S3 Table). In these experiments the control node in question was set at 1 while the rest of the control nodes varied. Then simulations were conducted using all possible initial conditions and the average proliferation activity was calculated. Constitutive activity of each hormone (=1) leads to higher 
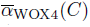
 values relative to the situation when the hormone is inactive (equal 0; Fig 4A). Statistical analysis revealed that under conditions used, CK, ETHL, and TDIF positively correlated with 
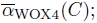
 and CK, IAA, BR, ETHL, and TDIF positively correlated with 
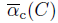
 at significance level 5% (Table 1). This outcome seems inconsistent with published experimental data on the importance of all hormones in cambium activity. However, as cambium tissue identity is maintained by TDIF/PXY signaling module, all hormonal signaling pathways most likely always incorporate this module. Therefore correlation between 
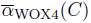
 and individual signals should be assessed while TDIF=1. Indeed, we found that all hormonal signals correlated with 
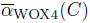
 or 
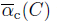
 under these conditions. Moreover, TDIF could amplify the effect of 391 each hormone on 
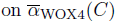
 (Fig 4B).

**Fig 4.**
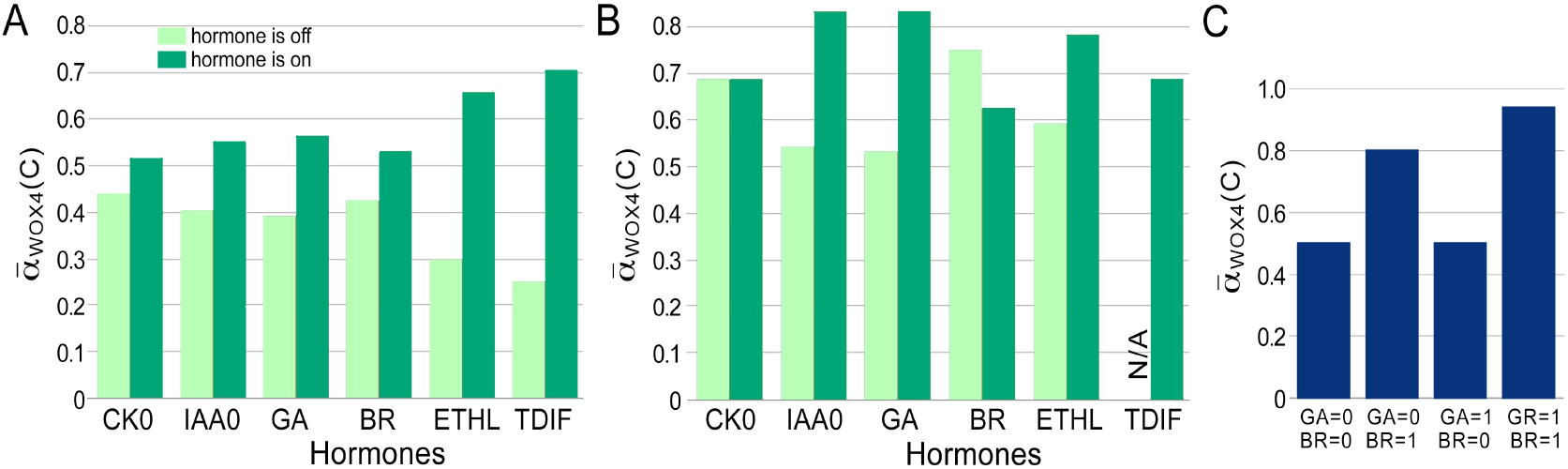
Effect of hormones on CARENET activity. A, 
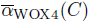
 obtained in the simulations experiments where corresponding hormone (control node) is turned on (dark green bars) or off (pale green bars). B, 
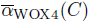
 obtained in the simulations experiments where TDIF was always on (TDIF=1) and corresponding hormone is turned on (dark green bars) or off (pale green bars). N/A, not applicable. C, Effect of gibberellins and brassinosteroids on 
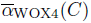
 under conditions when the rest of the control nodes were on.

Next, we analyzed relationships between GA and and BR. It appears that GA could

Increase 
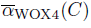
 to almost maximal value (1) in cooperation with BR, but not on its own (Fig 4C). These findings demonstrate synergistic relationships between gibberellic 3acid and brassinosteroids in promoting cambium proliferation.

To evaluate responsiveness of our system to changes in hormonal signals, we arbitrarily subdivided 
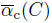
 into four bins: no activity 
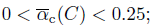
 low activity 
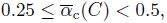
 medium activity 
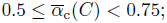
 and high activity 
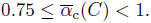
 With that calibration, medium activity was detected in 30 out of 64 possible control states, while high, low, and no activity was detected in 5, 4, and 25 control states respectively (S3 Table). 
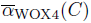
 behaved similarly achieving high, medium, low, and no activity in 30, 4, 0, and 30 control states respectively (S3 Table). Underrepresentation of the control states resulting in the low activity suggests that response of CARENET to hormonal signals is based on all-or-none principle. Once proliferation is activated, the role of hormones would be modulating the relative activity according to the developmental or environmental situation.

The fact that different combinations of control nodes result in similarly high or medium 
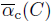
 values, emphasizes the redundancy of the hormonal signals in regulation of cambium activity. This situation is exemplified in Fig 3B, *C* where two different control states result in almost identical 
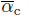
 values in structurally distinct final states. Furthermore, a given control state can also result in distinct final states with similar values of 
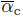
 for different initial conditions (S4 Fig). In the context of plant development these data suggest molecular heterogeneity of individual cells within the cambium.

### Testing the effect of mutations

Sensitivity of CARENET to hormonal signals suggests that it can be used to predict the impact of mutations on cambium proliferation. To test this hypothesis, we simulated the effect of mutations known to reduce cambium proliferation: *pxy* knockout [10,12]; double knockout of ERFs *erf108erf109* [24]; a quadruple knockout of IPT which was deficient in cytokinin synthesis [40]; and mutants in cytokinin receptor AHK4 [5,38]. Mutations were mimicked by keeping the values of PXY, ERF, IPT or AHK at 0 throughout the simulations. The 
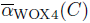
 failed to respond to TDIF in *pxy*, ethylene in *erf*, and to cytokinin in *ahp* and *ipt* (compare Fig 4A and Fig 5A–D). However, proliferation activity in these mutants responded to other signals. It demonstrates that corresponding nodes define sensitivity of the CARENET to specific external signals. In agreement with the experimental data, knockout of *PXY, ERF, AHK* and *IPT* resulted in lower proliferation activity (Fig 5F). These simulations demonstrate that CARENET represents experimental data on the functions of key signaling nodes in regulation of cambium proliferation. Next, we tested our hypothesis that BR controls activity of PXY/WOX4 module through WRKY by simulating *wrky* mutant. In agreement with our hypothesis *wrky* was insensitive to BR (Fig 5E) and exhibited lower proliferation activity than the control (Fig 5F).

**Fig 5.**
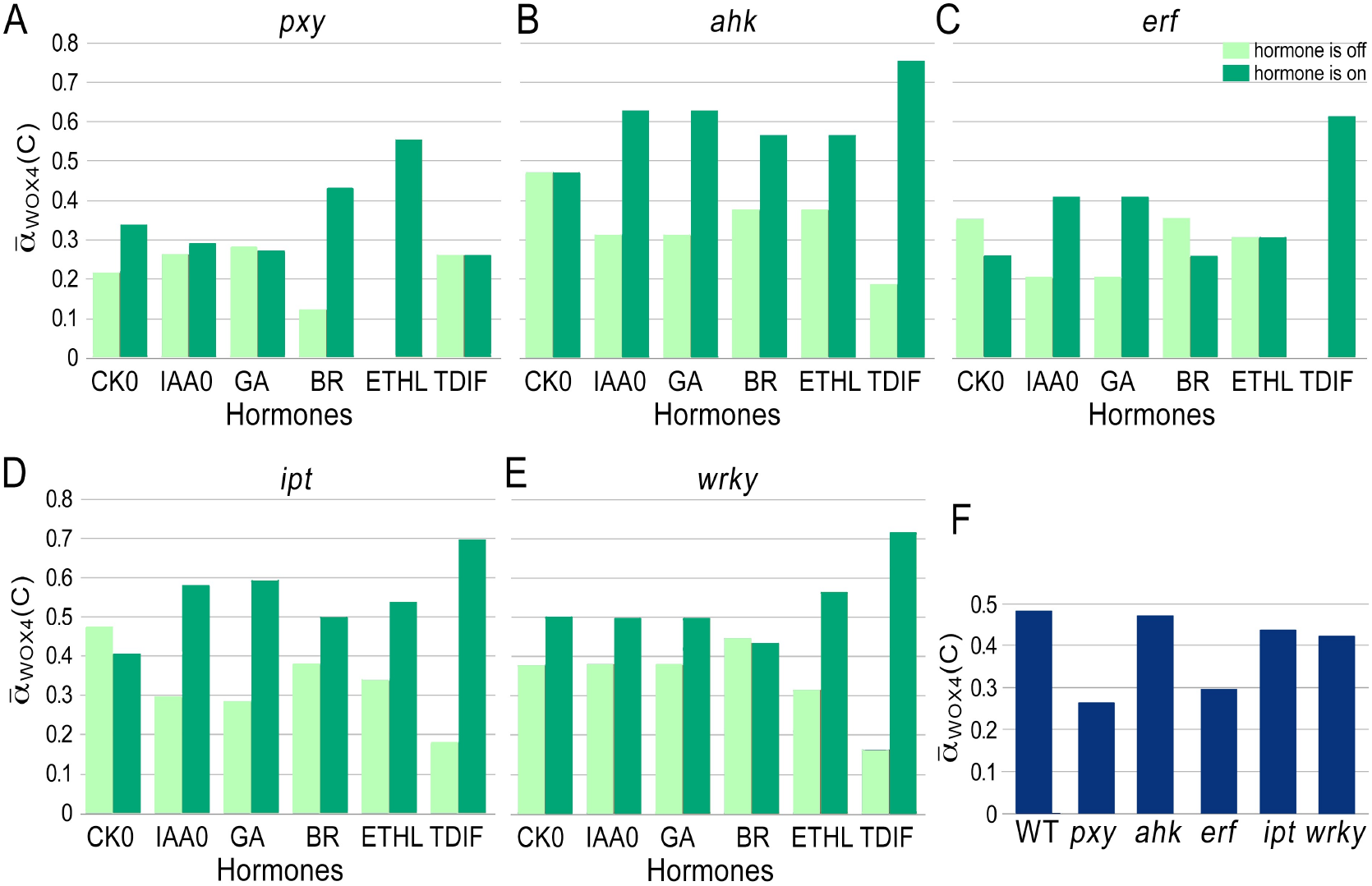
Effect of mutations on CARENET activity. A-E, Simulating effect of hormones on
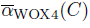
in *pxy*, *ahk*, *erf*, *ipt*, and *wrky* mutant backgrounds. 
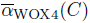
 was averaged for all the simulations with the corresponding control node on (dark green bar) or off (pale green bar). F, Proliferation in *pxy*, *ahk*, *erf*, *ipt*, and *wrky* mutant backgrounds.

## Discussion

### Hormonal cross talk in cambium regulation

The progress in understanding molecular mechanisms underlying cambium activity has benefited from analysis of procambium formation and vascular tissue patterning in *Arabidopsis* during embryogenesis, early post-embryonic growth [74,75], and secondary growth [2,23]. These efforts resulted in discovery of a number of genes responsible for each process. However, as mutations affecting procambium formation would also impact later cambium functions, the relevance of various sets of data for modeling cambium regulation requires further clarification [76]. In the absence of unequivocal experimental evidence to the contrary, it is reasonable to assume that these networks act in concert. Our computational experiments support the view that genetic mechanisms of procambium formation are likely to be exploited during secondary growth.

Our model combines the core cytokinin and auxin signaling modules employed by Benitez and Hejatko [20] with available experimental data on signaling mechanisms of ethylene, gibberellins, and brassinosteroids. One of the main challenges in this task was limited mechanistic data on the interaction between hormonal signals and the PXY/WOX4 module. We addressed this challenge by incorporating several nodes and edges that link distinct signaling pathways.

The first important integrating component of the CARENET is represented by the key ethylene signaling component ERFs. Microarray and qRT-PCR analyses demonstrated significant up-regulation of *ERFs* in *pxy* mutant background [24]. Cambium proliferation phenotype in *pxy* and *wox4* alleles was mild suggesting that up-regulation of the ethylene pathway compensates for inactivity of the PXY/WOX4 module. However, reduction of *WOX4* transcription in *erf109erf018* double mutant was statistically insignificant [24] indicating that more than two members of *ERF* gene family are responsible for this regulation. We anticipate that analysis of high-order *ERF* mutants would demonstrate regulation of *WOX4* or its close homologue *WOX14* by ERFs.

In addition to controlling *WOX4* transcription, ERFs link PXY/WOX4 module with cytokinin signaling. Although the effect of cytokinin on *WOX4* transcription has not been examined, RRB2 can bind to the promoter of *ERF1* and stimulate its transcription [43]. Correspondingly, transcription of *ERF1* was down-regulated in RRB mutant allele *arr2* [43]. Higher transcription level of *ERF* in combination with ethylene would stimulate transcription of *WOX4* and result in higher cambium activity. Consistent with this prediction, *AHK4* knockout allele with reduced sensitivity to cytokinin exhibits lower cambium activity [5]. ERF could also serve as a link between PXY/WOX4 module and BR signaling. This hypothesis has been supported indirectly by the fact that *wrky12* knockdown results in transcriptional down-regulation of ERF [67]. More experimental data is needed to clarity the role of WRKY in the regulation of cambium activity.

Linking ETHL, BR, GA, and TDIF with the *bona fide* components of auxin and cytokinin signaling produced a network that accurately represents experimental data in showing additive effect of hormones on cambium activity and mimicking the effect of mutations in the key nodes. More nodes and edges can be added to the CARENET to facilitate understanding of how cambium proliferation changes in response to other hormones, mobile peptides, RNAs, environmental changes and nutrient availability.

### Molecular heterogeneity of cambium cells

Being a typical Boolean network, CARENET lacks physical time scales. Therefore the number and structure of steps preceding establishment of the stable states or limit cycles shown in Fig 3, and S1-S4 Figs lack biological relevance. These steps depend on the structure of the network and the rules. Likewise, the number and structure of steps in the limit cycles are time-independent. However, activity of specific nodes in the limit cycles (i.e. the ratio of the number of steps at which a node acquires value 1 to the number of all steps in a cycle) is useful for converting the binary readouts of a Boolean network into a continuous scale. In turn, a continuous scale enables comparison of gene activity in distinct final conditions generated by different combinations of controls. Consequently, results of the simulations could be interpreted more accurately in a specific biological context.

Our simulations demonstrate that proliferation of cambium is controlled redundantly by hormonal signals (control nodes). For example the combination of CK0 and ETHL produced the same proliferation activity (0.7) as GA and TDIF (S3 Table). The effect of each signaling pathway appears to be additive and higher proliferation could be achieved when more pathways become active. This outcome supports available experimental evidence on the redundancy of mechanisms regulating developmental processes. Less expectedly, our simulations showed that identical control states may result in structurally distinct stable states with similar proliferation activity 
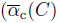
 or
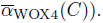
. Thus, individual cambium cells exhibiting the same proliferation activity may possess distinct molecular identities. Such identities can be characterized by the different expression levels of the key regulatory genes. While activity of some pathways in the individual cells could be suboptimal, the proliferation would still be sustained by elevated activity of other pathways.

Temporal transcriptional analysis demonstrated that successive stages of lateral root meristem establishment are accompanied by sequential activation of different sets of genes [77]. Similar processes may take place during the establishment of all meristems. Nonetheless, subsequent activity of the stem cell niches including cambium and apical meristems can potentially undergo stochastic oscillations. Predicted molecular heterogeneity of cambium cells could also occur in other meristems and it would be interesting to verify this hypothesis using single-cell transcriptomics.

## Limitations and future work

1. Currently the only receptor kinase in the CARENET is PXY. The role of PXY in the regulation of cambium proliferation was supported by the following observations: (i) reduction of cambium cell niche in *pxy/tdr1* mutants [10, 12]; and (ii) number of procambium cells in stems increases following treatment with TDIF [10]. However, proliferation of cambium is not abrogated in *pxy* suggesting existence of functionally redundant signaling mechanisms. Recently, other receptor kinases were implicated in the regulation of cambium proliferation, e.g. *PXY* correlated PXC [78] and REDUCED IN LATERAL GROWTH1 [79]. It is also plausible that cambium is regulated by as yet uncharacterized signaling modules. The limited experimental data on these signaling processes precludes adding them to CARENET at present. It would be desirable to incorporate these nodes as more information becomes available.
2. Many of the nodes in CARENET represent not a single gene, but a gene family with overlapping expression pattern, e.g. *ARF, RRA, WRKY, ERF*. Some members of these families may even have antagonistic functions. For example whilst AHPs are generally positive regulators of CK response, AHP6 inhibits CK signaling [39]. Functional redundancy could also confound determining the function and downstream targets of some nodes. The CARENET could be further refined once functions of the members of gene families are determined experimentally.
3. Our current model takes into account only external GA; however intracellular homeostasis of GA could also be important for cambium activity. The importance of the internally produced GA on secondary growth remains unknown. Filling this knowledge gap would establish the feedback loop for GA and introduce additional edges connecting GA loop to other parts of the network.

## Conclusions

This work aims at understanding regulation of cambium proliferation by hormonal signals. Addressing this problem experimentally is challenging because of pleiotropic effect of hormones on gene transcription. Appropriately selected mathematical modeling tools can facilitate both experimental design and interpretation of the experimental data. While analytical solutions can be effective for dealing with simple processes, complex multi-parametric phenomena are more efficiently addressed using simulations. We have constructed and computationally verified a Boolean deterministic network model for the regulation of cambium activity. Behavior of the CARENET corroborates published experimental data.

To quantify behavior of CARENET under different combinations of hormones, we have introduced and compared the average activity coefficients of genetic markers *WOX4* and *ATHB8* whose levels of expression are known to correlate strongly with proliferation. The average activity coefficients were computed using a method inspired by classical statistical mechanics. The central idea of the method is to use ensemble averaging over an ensemble consisting of all final states attainable with a particular control state. In statistical mechanics, every initial condition is assigned a weight (the probability of occurrence of that condition), and then quantities of interest are averaged over the ensemble of all relevant initial conditions using the above probabilities as weights. In the present case, we propose to assign weights to each *final* state based on the number of different initial conditions leading to that state. Then, given a specific control state, the average activity of the markers is computed by (i) calculating their activity for each final state, and (ii) averaging these activities over the ensemble of the final states using the above assigned weights. The proposed ensemble averaging procedure is applicable to generic Boolean networks, and thus it may be of broad interest in computational biology.

Overall, our network accurately represents mutually inhibitory relationships between auxin and cytokinin as well as cooperation between gibberellic acid and brassinosteroids during cambium proliferation. CARENET can be used for predicting: (i) cambial activity in response to developmental cues; (ii) effect of mutations on gene transcription and cambium proliferation; (iii) co-expression of genes and genetic markers. In addition to resembling available experimental data, CARENET demonstrates that similar cambium proliferation activity can be achieved under multiple final states. Hence, individual cells in cambium and potentially other stem cell niches can have distinct molecular identities. Another important feature of CARENET is that it can be expanded to include components of other signaling pathways and ultimately be integrated with existing models of xylem and phloem differentiation to compose a comprehensive model of secondary growth. Such a model would enable predicting the key nodes that can be targeted in plant breeding programs for optimized yield and biomass quality. In particular, breeding trees with higher wood yield and more efficient sequestration of CO_2_ from the atmosphere would both alleviate the greenhouse effect and augment wood production.

## Acknowledgments

The authors are grateful to Deirdre Fahy for thoughtful notes and for proofreading the text. This project was supported by NIFA hatch project WNP00826 (to AS).

## Supporting Information

**S1 Fig. Simulation of the CARENET which results in a stable state at step 26 shown in Figure 3A. Control nodes CK0, IAA0, EHTL and TDIF are on.**

**S2 Fig. An example of limit cycle where cytokinin (CK0), auxin (IAA0), and brassinosteroids are on**. The cycle of 6 steps long establishes at step 9. Brackets indicate single cycles.

**S3 Fig. An example of limit cycle where cytokinin (CK0), auxin (IAA0), and brassinosteroids are on.** The cycle of 6 steps long establishes at step 9. Brackets indicate single cycles.

**S4 Fig. Two structurally distinct limit cycles with high proliferation activity (
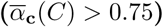
) generated by identical set of controls.**

**S1 Table. Rules for calculating the status of each node in the CARENET.** The rules are inferred form the experimental evidence shown in S2 Table.

**S2 Table. Experimental evidence supporting interactions between nodes of CARENET.**

**S3 Table. Statistical analysis of all stable states generated by the CARENET.**

